# Host phenology regulates ecological and eco-evolutionary feedbacks underlying parasite-host demographic cycles

**DOI:** 10.1101/2021.06.07.447391

**Authors:** Hannelore MacDonald, Dustin Brisson

## Abstract

Parasite-host interactions can result in periodic population dynamics when parasites over-exploit host populations. The timing of host seasonal activity, or host phenology, determines the frequency and demographic impact of parasite-host interactions which may govern if the parasite can sufficiently over-exploit their hosts to drive population cycles. We describe a mathematical model of a monocyclic, obligate-killer parasite system with seasonal host activity to investigate the consequences of host phenology on host-parasite dynamics. The results suggest that parasites can reach the densities necessary to destabilize host dynamics and drive cycling in only some phenological scenarios, such as environments with short seasons and synchronous host emergence. Further, only parasite lineages that are sufficiently adapted to phenological scenarios with short seasons and synchronous host emergence can achieve the densities necessary to over-exploit hosts and produce population cycles. Host-parasite cycles can also generate an eco-evolutionary feedback that slows parasite adaptation to the phenological environment as rare advantageous phenotypes are driven to extinction when introduced in phases of the cycle where host populations are small and parasite populations are large. The results demonstrate that seasonal environments can drive population cycling in a restricted set of phenological patterns and provides further evidence that the rate of adaptive evolution depends on underlying ecological dynamics.

## 1 Introduction

The impact of inter-species interactions on population demography is a function of both the abundance and activity patterns of the interacting species. For example, the abundance of both a predator and prey species determines the prevalence of predation which, in turn, alters the demographic dynamics of one or both species. Some ecological interactions have even been shown to drive population sizes to fluctuate cyclically over time.^1^ Ecological interactions leading to population cycles have been studied extensively in predator-prey, herbivore-plant, and parasite-host systems, although the importance of seasonal activity patterns is rarely considered. Seasonal activity patterns determine the temporal abundance of a population which modifies the strength of inter-species interactions.^2–6^ Here we explore the consequence of seasonal activity patterns on parasite-host population dynamics and how seasonal patterns can result in parasite-host population cycles. Additionally, we explore how parasite-host population cycles can alter the rate of parasite virulence evolution.

Population cycling generally starts with an over-exploitation of resources followed by a population crash that allows resources to rebound.^7^ In the classic lemming demographic cycles, lemmings over-consume plant resources resulting in dramatic declines in lemming population sizes in subsequent years due to plant scarcity.^8^ The plant populations are released from lemming herbivory and increase in abundance, providing sufficient resources for lemming population growth and a restart of the demographic cycle. Intrinsic, delayed density dependent drivers such as these can account for the periodic or quasi-periodic oscillatory population dynamics observed in many ecologically coupled systems.^1^

Seasonal activity patterns, or phenology, influence the impact of inter-species interactions on demographic dynamics.^9,10^ That is, seasonal activity patterns determine the number and type of inter-species interactions by altering the proportion of a population that is active throughout the year. Prior theoretical research demonstrated that the total number of parasite infections, which determines the parasite population size, varied dramatically among the host phenological patterns explored.^11^ Further, the virulence strategies that maximize parasite fitness also differed among phenological patterns due to the differences in the temporal distribution of new infections. However, this work restricted host demographic dynamics such that the potential for population cycles subsequent effect on evolutionary dynamics could not be investigated.

Changes in the population sizes of interacting species that result from ecological interactions can also influence the rate or direction of evolutionary change.^12^ These eco-evolutionary feedbacks arise when evolutionary change occurs on time-scales congruent with ecological change. For example, evolutionary adaptation of parasites to a specific host phenological pattern increases parasite densities with a concomitant decrease in host population sizes, which alters both the ecological interactions and the strength and direction of natural selection. Increases in parasite fitness could result in a parasite population that can over-exploit hosts leading to temporal oscillations in population sizes with concomitant oscillations in infection prevalence and the strength of natural selection. Thus, host phenology could create conditions that drive the evolution of sufficiently high parasite densities to destabilize host populations and drive population cycles. The resulting population cycles, in turn, could influence the rate and direction of further evolutionary change.

Here we explore eco-evolutionary feedbacks driven by parasite infection in a seasonal environment. We extend a previously published modeling framework^11^ to follow within-season transmission dynamics as well as between-season parasite and host demography to determine if evolutionary increases in parasite fitness can lead to periodic population dynamics given different host phenological patterns. Further, we investigate how changes in parasite and host demography, including population cycling, can influence the rate and direction of parasite evolution in seasonal environments. These results contribute to the longstanding goal of revealing how cycling arises by showing how ecological and evolutionary interactions can generate periodic dynamics.

## 2 Model description

The model describes the transmission dynamics of a free-living, obligate-killer parasite that infects a seasonally available host (Figure 1). The size of the emerging host cohort in season *n, ŝ*(*n*), is determined by the number of hosts that reproduced in season *n −* 1. *ŝ*(*n*) enter the system at the beginning of the season over a period given by the function *g*(*t, t*_*l*_). Hosts, *s*, have non-overlapping generations and are alive for one season. The parasite, *v*, infects hosts while they are briefly susceptible early in their development (*e*.*g*. baculoviruses of forest *Lepidoptera*^13–17^ and univoltine insects parasitized by ichneumonid wasps^18–20^). The parasite must kill the host to release new infectious progeny. The parasite completes one round of infection per season because the incubation period of the parasite is longer than the duration of time the host spends in the susceptible developmental stage. This transmission scenario occurs in nature if all susceptible host stages emerge over a short period of time each season so that there are no susceptible host stages available when the parasite eventually kills its host. Parasites may also effectively complete only one round of infection per season if the second generation of parasites do not have enough time in the season to complete their life cycle in the short-lived host.

**Figure 1:**
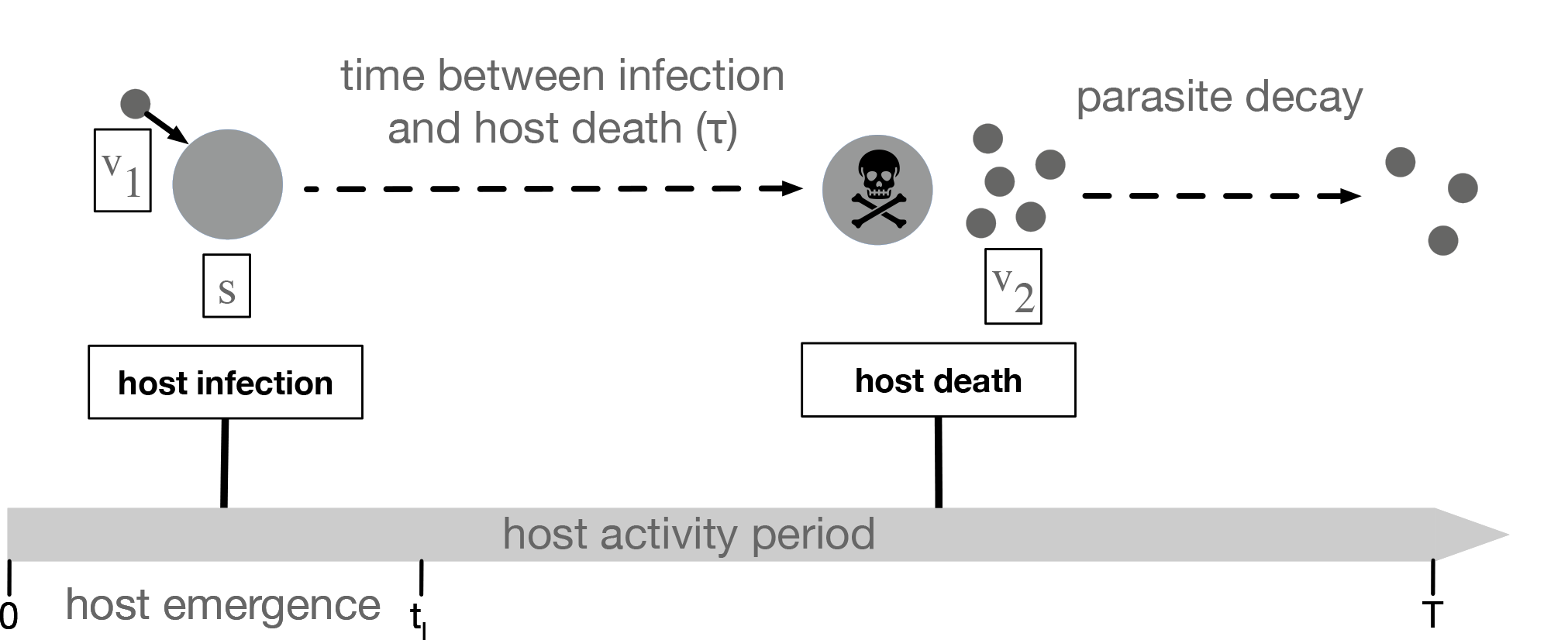
Diagrammatic representation of the infectious cycle within each season. The host population (*ŝ*(*n*)) at the start of season *n* are the offspring of uninfected hosts that survived and reproduced at the end of the prior season. The parasite population at the start of season 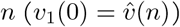 are derived from infected hosts killed by the parasite prior to the end of season *n* 1 and survived in the environment until the end of the season *T* (*v*_2_(*T*)). All parasites emerge at the beginning of the season (*t* = 0) while all hosts emerge at a constant rate between time *t* = 0 and *t* = *t*_*l*_. The rate of new infections is density dependent resulting in the majority of infections occurring near the beginning of the season when susceptible host and free parasite densities are high. Parasite-induced host death at time *τ* post-infection releases parasite progeny (*v*_2_) into the environment where they decay in the environment at rate *δ*. The monocyclic parasite progeny, *v*_2_, do not infect uninfected hosts within the same season. Parasite progeny that survive in the environment to the end of the season comprise the parasite population that emerge in the following season 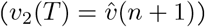.

We ignore the progression of the susceptible stage, *s*, to later life stages as it does not impact transmission dynamics. To keep track of these dynamics, we refer to the generation of parasites that infect hosts in the beginning of the season as *v*_1_ and the generation of parasites released from infected hosts upon parasite-induced death as *v*_2_. The initial conditions in the beginning of the season are thus 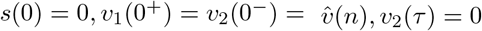, where 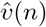 is the size of the starting parasite population introduced at the beginning of season *n* determined by the number of parasites produced in season *n −* 1. The transmission dynamics in season *n* are given by the following system of delay differential equations (all parameters are described in Table 1):

**Table 1:** Model parameters and their respective values.

| Parameter | Description | Value |
| --- | --- | --- |
| $s$ | susceptible hosts | state variable |
| $v_1$ | parasites that infect hosts in current season | state variable |
| $v_2$ | parasite progeny released in current season | state variable |
| $\hat{v}(n)$ | starting parasite population in season $n$ | state variable |
| $\hat{s}(n)$ | host cohort in season $n$ | state variable |
| $t_l$ | length of host emergence period | time (varies) |
| $T$ | season length | time (varies) |
| $\alpha$ | transmission rate | $3.5 * 10^{-7}/(\text{parasite} \times \text{time})$ |
| $\beta$ | number of parasites produced upon host death | 200 parasites |
| $\delta$ | parasite decay rate in the environment | 2 parasites/parasite/time |
| $d$ | host death rate | 0.25 hosts/host/time |
| $\tau$ | time between host infection and host death (1/virulence) | time (evolves) |
| $\sigma$ | host fecundity | 500 hosts |
| $\rho$ | density dependent parameter | 0.0001 |

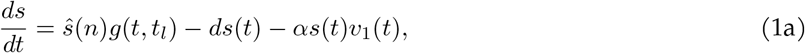

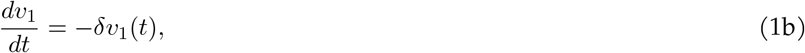

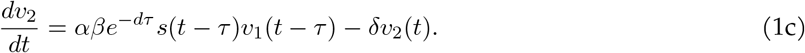

where *d* is the host death rate, *δ* is the decay rate of parasites in the environment, *α* is the transmission rate, *β* is the number of parasites produced upon host death and *τ* is the delay between host infection and host death. *τ* is equivalent to virulence where low virulence parasites have long *τ* and high virulence parasites have short *τ*. We make the common assumption for free-living parasites that the removal of parasites through transmission (*α*) is negligible,^17, 21, 22^ *i*.*e*. (1b) ignores the term *−αs*(*t*)*v*_1_(*t*).

The function *g*(*t, t*_*l*_) is a probability density function that captures the per-capita host emergence rate by specifying the timing and length of host emergence. We use a uniform distribution (*U* (*•*)) for analytical tractability, but other distributions can be used.

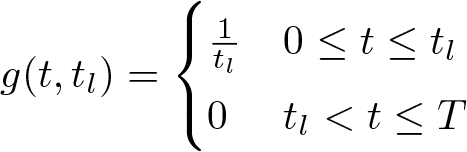

*t*_*l*_ denotes the length of the host emergence period and *T* denotes the season length. The season begins (*t*_0_ = 0) with the emergence of the susceptible host cohort, *ŝ*. The host cohort emerges from 0 *≤ t ≤ t*_*l*_. *v*_2_ parasites remaining in the system at *t* = *T* give rise to next season’s initial parasite population 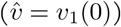. Parasites that have not killed their host by the end of the season do not release progeny. Background mortality arises from predation or some other natural cause. We assume that infected hosts that die from background mortality do not release parasites because the parasites are either consumed or the latency period corresponds to the time necessary to develop viable progeny.^23, 24^ We solve equations 1a-c analytically Appendix A.

### 2.0.1 Between-season dynamics

To study the impact of the feedback between host demography and parasite fitness on parasite evolution we let the size of the emerging host cohort be a function of the number of uninfected hosts remaining at the end of the prior season

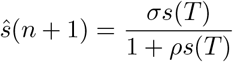

where *σ* is host reproduction and *ρ* is the density dependent parameter.

Host carryover generates a feedback between parasite fitness and host demography that drives cycling for some parameter ranges. We primarily explore the between-season dynamical behavior of the model numerically. We discuss the stability analysis in more detail in Appendix A.

### 2.0.2 Parasite fitness

A parasite introduced into a naive host population persists or goes extinct depending on the length of the host emergence period and season length. The stability of the parasite-free equilibrium is determined by the production of *v*_2_ resulting from infection of *s* given by

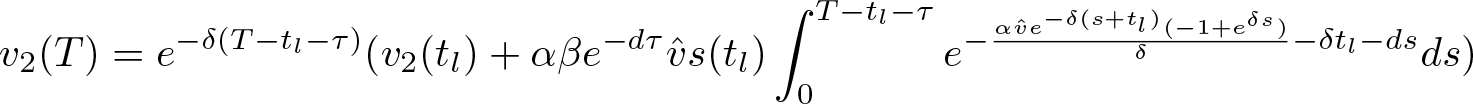

The parasite-free equilibrium is unstable and a single parasite introduced into the system at the beginning of the season will persist if the density of *v*_2_ produced by time *T* is greater than or equal to 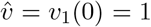 (*i*.*e. v*_2_(*T*) *≥* 1, modulus is greater than unity). This expression is a measure of a parasite’s fitness when rare given different host phenological patterns. See Appendix A for details of the analytical solution.

### 2.0.3 Parasite evolution

To study how parasite traits adapt given different seasonal host activity patterns, we use evolutionary invasion analysis.^25, 26^ We first extend system (1) to follow the invasion dynamics a rare mutant parasite

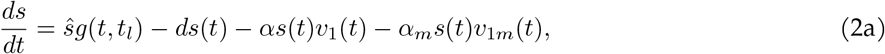

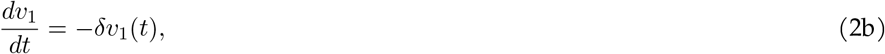

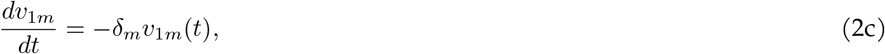

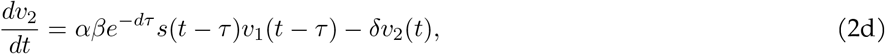

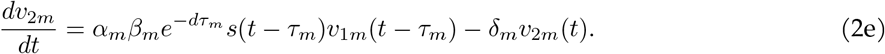

where *m* subscripts refer to the invading mutant parasite and its corresponding traits. See Appendix B for details of the time-dependent solutions for equations (2a-2e).

The invasion fitness of a rare mutant parasite depends on the density of *v*_2*m*_ produced by the end of the season (*v*_2*m*_(*T*)) in the environment set by the resident parasite at equilibrium density 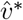. The mutant parasite invades in a given host phenological scenario if the density of *v*_2*m*_ produced by time *T* is greater than or equal to the initial *v*_1*m*_(0) = 1 introduced at the start of the season (*v*_2*m*_(*T*) *≥* 1).

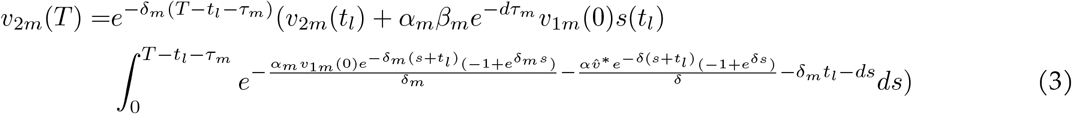

To study the evolution of virulence traits, we first assume all other resident and mutant traits are identical (*e*.*g. α* = *α*_*m*_). Note that because there is no trade-off between *β* and *τ*, the parasite growth rate in the host is essentially the trait under selection. That is, *β* is constant regardless of *τ* thus the time between infection and when the parasite kills the host and releases new parasites is the rate that *β* new parasites are assembled. To find optimal virulence for a given host phenological scenario, we find the uninvadable trait value that maximizes (3). That is, the virulence trait, *τ* ^*∗*^, that satisfies

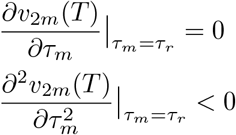

Note that the measure in equation (3) incorporates the effect of the resident on the population state (number of susceptibles over one season), which means that it is not a measure of *R*_0_ (which by definition assumes a non-disease environment). Thus, we can use *v*_2*m*_(*T*) as defined in (3) as a maximand in evolutionary dynamics.^27^

We previously found the the uninvadable virulence trait value, *τ* ^*∗*^ that maximizes (3) for system with a constant host cohort size each season.^11^ In this study we found that an intermediate *τ* ^*∗*^ maximizes parasite fitness and that the exact host phenology pattern determines optimal *τ* ^*∗*^. In the present study, host carryover from one season to the next creates a feedback between parasite fitness and host demography that drives cycling for some parameter ranges. When parasite-host dynamics are cycling, (3) no longer reliably predicts the outcome of parasite evolution. We thus conduct simulation analysis to verify that the evolutionary stable level of virulence is qualitatively the same as previous results.

The simulation analysis was done by first numerically simulating system (1) with a monomorphic parasite population. A single mutant parasite is introduced at the beginning of the season after 100 seasons have passed. The mutant’s virulence strategy is drawn from a normal distribution whose mean is the value of *τ* from the resident strain (*τ*_*m*_ = *τ*_*r*_ + *𝒩* (0, 0.1)). System (2) is then numerically simulated with the resident and mutant parasite. New mutants arise randomly after 1000 seasons have passed since the last mutant was introduced, at which point system (2) expands to follow the dynamics of the new parasites strain. This new mutant has a virulence strategy drawn from a normal distribution whose mean is the value of *τ* from whichever parasite strain has the highest density. System (2) continues to expand for each new mutant randomly introduced after at least 1000 seasons have passed. Any parasite whose density falls below 1 is considered extinct and is eliminated. Virulence evolves as the population of parasites with the adaptive strategy eventually invade and rise in density. Note that our simulations deviate from the adaptive dynamics literature in that new mutants can be introduced before earlier mutants have replaced the previous resident. Previous studies have shown that this approach is well suited to predicting evolutionary outcomes.^28–30^

## 3 Results

Parasites with high fitness in some seasonal environments can drive dynamic parasite-host cycles which resemble classic consumer-resource cycles (Figure 2). In the present model, parasites that can achieve sufficiently high densities infect and sterilize a substantial proportion of the univoltine host population resulting in both a decrease in the host population size and an increase in the parasite population size in subsequent years. The resulting small host populations sizes limit infections which leads to a dramatic decrease in parasite population size in the following years. Very small parasite populations, in turn, release the host population from parasite-mediated density control allowing the host population to increase in size. This cycle continues with large host populations supporting rapid parasite population growth which then drives down the size of the host population. In the current model, one complete cycle requires at least four years with parasite population size peaks trailing the host population peaks by two to three years.

**Figure 2:**
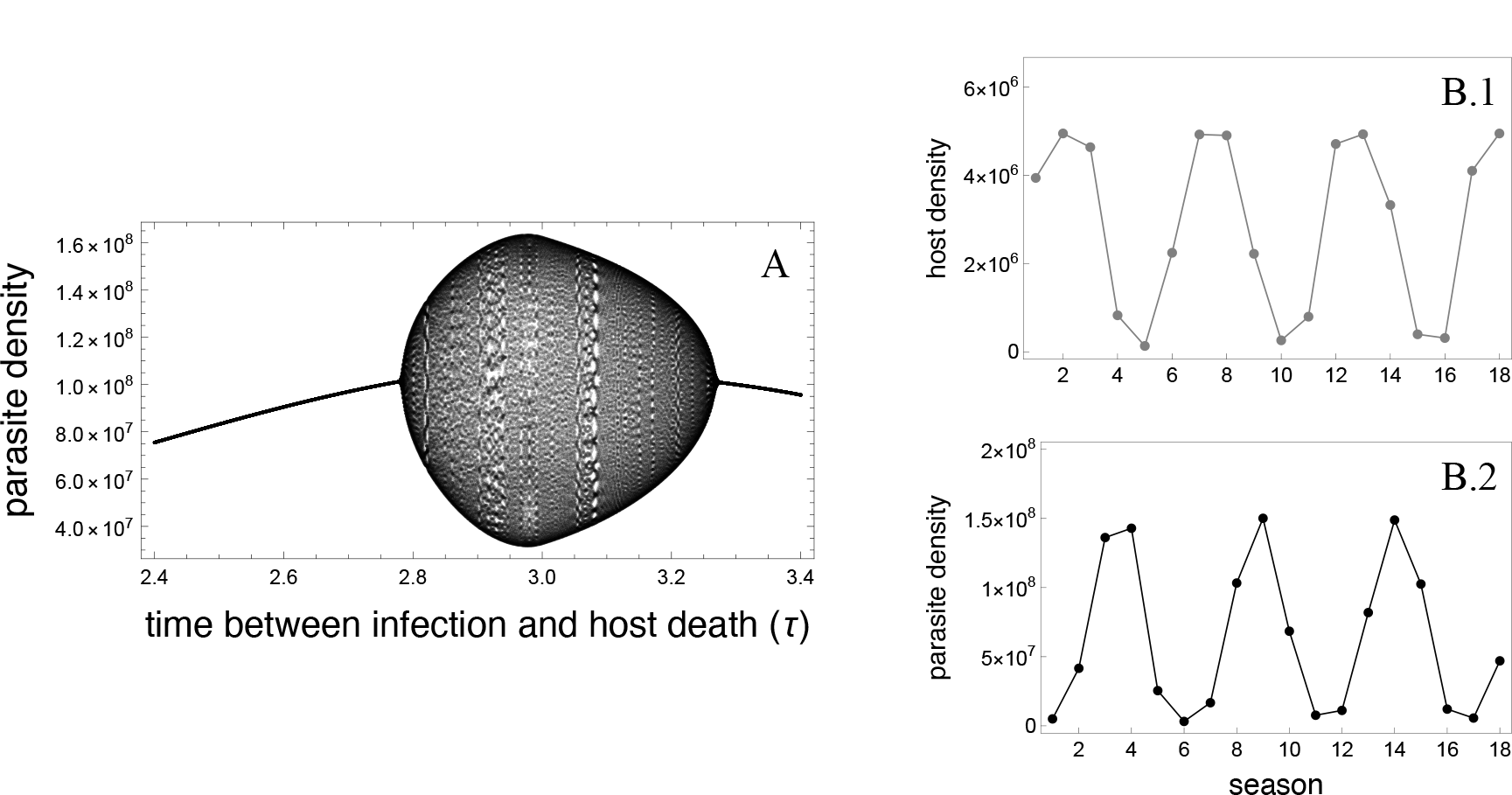
High-fitness parasites can drive multi-season epidemic cycles. **A**. Parasite density increases as the virulence phenotype approaches the time between infection and host death (*τ*) that maximizes parasite fitness.^11^ Parasite populations can reach sufficiently high densities in some host phenological patterns, as seen in **A**, to destabilize demographic dynamics resulting in a bifurcation that drives quasiperiodic parasite-host dynamics. The bifurcation diagram shows end of season parasite densities for parasites with different virulence phenotypes (*τ*) during seasons 800-900 in a system where the host season is short (*T* = 4) and hosts emerge synchronously (*t*_*l*_ = 1). The most fit parasites (2.75 *< τ <* 3.26) achieve densities that can disrupt dynamics and cause cycling. Parasites with virulence phenotypes that are too high (*τ <* 2.75) or too low (*τ >* 3.26) do not cause parasite-host cycles in this host phenological environment. **B**. The population dynamics of hosts (**B.1**) and parasites (**B.2**) in a system experiencing quasiperiodic population cycles (*τ* = 2.8, *T* = 4, *t*_*l*_ = 1, other parameters found in Table 1) after reaching the quasiperiodic attractor. High parasite densities (ex. season 3-4) infect and sterilize a large proportion of the host population resulting in a dramatic host population decline (ex. seasons 4-5). The limited number of susceptible hosts causes a subsequent decline in parasite populations (ex. seasons 5-7). Host density rebounds once relieved from infection pressure (ex. seasons 6-8) allowing the parasites to exploit the host population again, driving a continuation of quasiperiodic cycling. In both panels: *T* = 4, *t*_*l*_ = 1, all other parameters found in Table 1.

Host phenological scenarios either do not cause inter-annual population cycling or cause quasi-periodic parasite-host population cycles (Figure 3). In the majority of scenarios in which cycling occurs, the discrete dynamics form a closed invariant curve in the phase plane in which the phase is incommensurate and thus the asymptotic trajectory fills the invariant curve by never repeating itself (Fig 5, left panel). That is, the population sizes for both the host and parasite do not repeat across seasons, resulting in quasi-periodic cycles that are likely generated by a Neimark-Sacker bifurcation (see Appendix A.)

**Figure 3:**
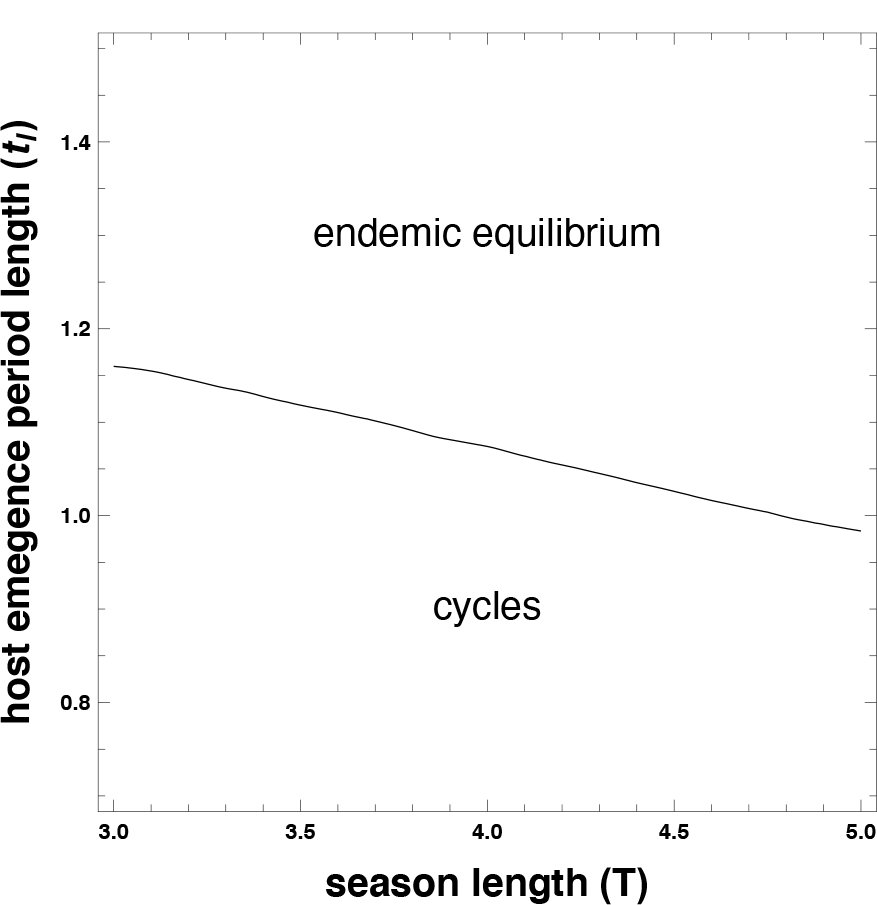
Parasite-host cycles occur in some, but not all, host phenological patterns. Boundary plot shows host phenological patterns where dynamics are stable (”endemic equilibrium”) or cycling for parasites possess the optimal virulence trait for their phenological environment. Parasites are more likely to achieve the densities necessary to drive cycles when host emergence periods are short (small values of *T*) and host emergence is synchronous (small values of *t*_*l*_). All other parameters found in Table 1.

Quasi-periodic parasite-host population cycles occur when hosts exhibit some, but not all, phenological patterns (Figure 3). Host phenological patterns influence parasite densities, and thus if cycling occurs, both due to parasite evolution and to ecological interactions. Evolutionarily, the level of virulence that maximizes parasite fitness, and thus maximizes the density of parasites, differs among host phenological patterns. Nevertheless, parasites with the virulence trait that optimizes fitness in some phenological environments do not reach densities sufficient to destabilize equilibrium dynamics and cause parasite-host population cycles. Generally, parasites adapted to environments with shorter host activity seasons can reach sufficiently high densities to provoke parasite-host population cycles. The short-season adapted parasites reach higher population sizes as fewer infected hosts die of natural causes before parasites can release their progeny. By contrast, a greater number of infected hosts die through natural host mortality when host activity seasons are longer, resulting in fewer infected hosts contributing parasite progeny to the following season.

Parasites adapted to phenological environments with limited variation in the time when each host first emerges within a season are more likely to achieve population cycle-inducing densities than environments with more variable host emergence timing (Figure 3). Synchronous host emergence results in all infections occurring near simultaneously such that parasites with the optimal virulence trait will kill all infected hosts near the end of the season in order to minimize decay of parasite progeny in the environment. By contrast, greater host emergence variability results in concomitant variability in the timing of infections. Parasites adapted to phenological environments with greater host emergence variation have a virulence level that causes hosts infected early in the season to release progeny too early -where parasites decay in the environment -and to not kill hosts infected later in the season where hosts die naturally without producing parasite progeny. The loss of potential parasite progeny through environmental decay or non-parasite induced death of infected hosts limits the parasite density to levels below those that can destabilize host-parasite dynamics and cause demographic cycles.

Conditions that promote high parasite density, such as low environmental decay, can also increase the parameter region where cycling occurs (Table 2). High parasite density results in highly synchronous infections early in the season which increases parasite density and the likelihood that parasites destabilize host dynamics. Conversely, conditions that prevent high parasite density, such as a high natural host mortality rate, decrease the parameter range where cycling occurs. A high host mortality rate increases the death rate of infected hosts and thus the number of infections that successfully release new parasites. When fewer infections release new parasites, the parasite population is less likely to reach densities that generate cycles.

**Table 2:** The impact of each variable on demographic cycles.

| Increases in variable value | Impact on possibility of demographic cycling |
| --- | --- |
| season length ($T$) | ↓ cycling |
| emergence period length ($t_l$) | ↓ cycling |
| host mortality ($d$) | ↓ cycling |
| decay rate ($\delta$) | ↓ cycling |
| transmission rate ($\alpha$) | ↑ cycling |
| parasites released at host death ($\beta$) | ↑ cycling |
| host fecundity ($\sigma$) | ↑ cycling |

Parasite-host population cycles impede the rate at which parasites adapt to host phenological environments (Figures 4, 5). Rare advantageous mutations readily invade systems in which the populations are not cycling. However, the phase of a population cycle at which a rare advantageous mutant is introduced into a system determines if it will displace the resident parasite despite their fitness advantage. Rare advantageous mutants invade cycling systems only in seasons when the resident parasite population is at low densities and the host population size is increasing or is at high densities. By contrast, advantageous mutants rarely find a susceptible host when the resident parasite is at high densities and the host population at low densities due to the numerical advantages of the resident. This eco-evolutionary feedback results in the loss of many advantageous mutations and a reduced rate of evolution toward the virulence strategy that optimizes parasite fitness.

**Figure 4:**
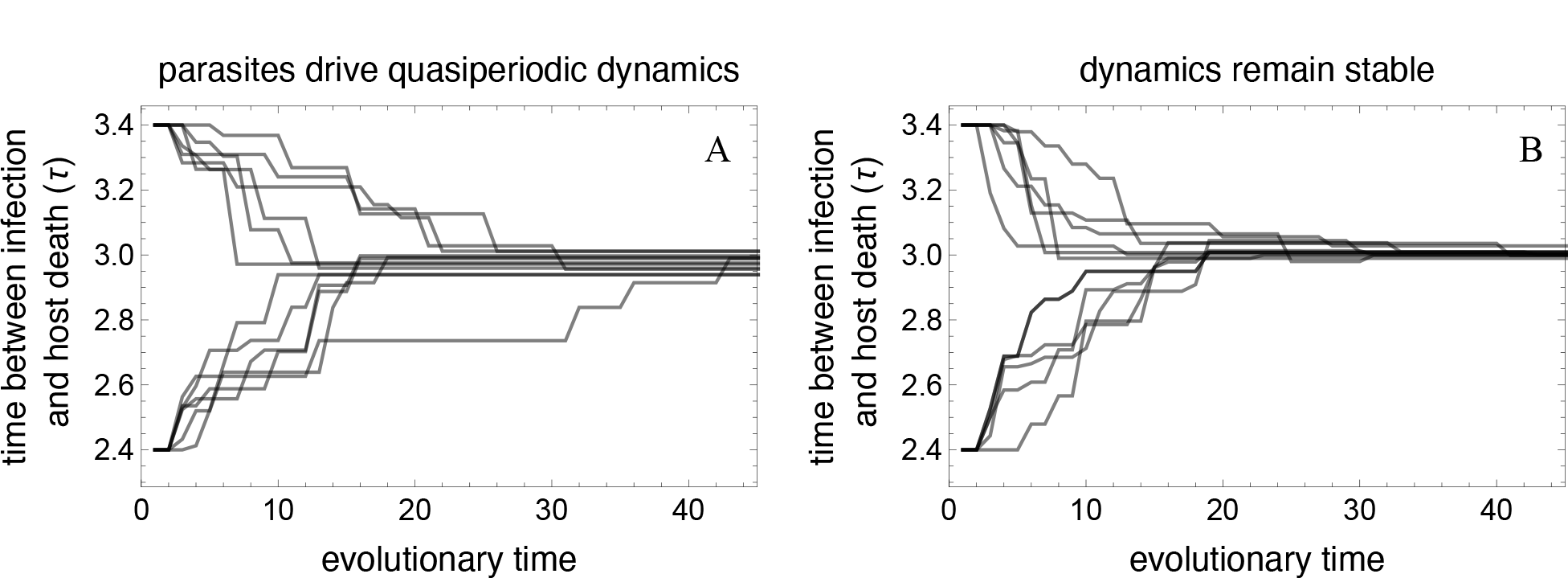
Adaptive evolution proceeds more slowly during parasite-host demographic cycles than in stable equilibrial systems. Parasite evolution toward an intermediate optimal virulence strategy occurs more slowly when population demography is cycling (**A**) than in an equilibrial dynamic system (**B**). (**A**) Increases in parasite density as parasites evolve drive demographic cycling for 2.75 *< τ <* 3.26. Population cycling delays adaptive evolution as rare mutants fail to invade the system when introduced at many phases of the dynamic cycle despite their selective advantage. By contrast, rare advantageous mutants always invade systems with stable dynamics (**B**). Plots show twelve independent simulations for each set of parameters -six runs starting at a virulence level lower than optimum and six runs starting at a virulence level higher than the optimum -where an adaptive mutant is introduced into the population no more than once every 1000 seasons. Evolutionary time represents the cumulative number of adaptive mutants sequentially introduced into each population. The average evolutionary time needed to reach the optimal virulence strategy is higher in the cycling system (**A**. 21 mutants, range: 6-42 mutants) than in the stable system (**B**. 14 mutants, range: 6-27 mutants) Population cycling could not occur in **B** as the host cohort size remained constant across seasons (*ŝ* = 10^8^); host cohort size in **A** (*ŝ*(*n*)) was determined by the number of hosts that reproduced in season *n −* 1. *T* = 4, *t*_*l*_ = 1, all other parameters found in Table 1.

**Figure 5:**
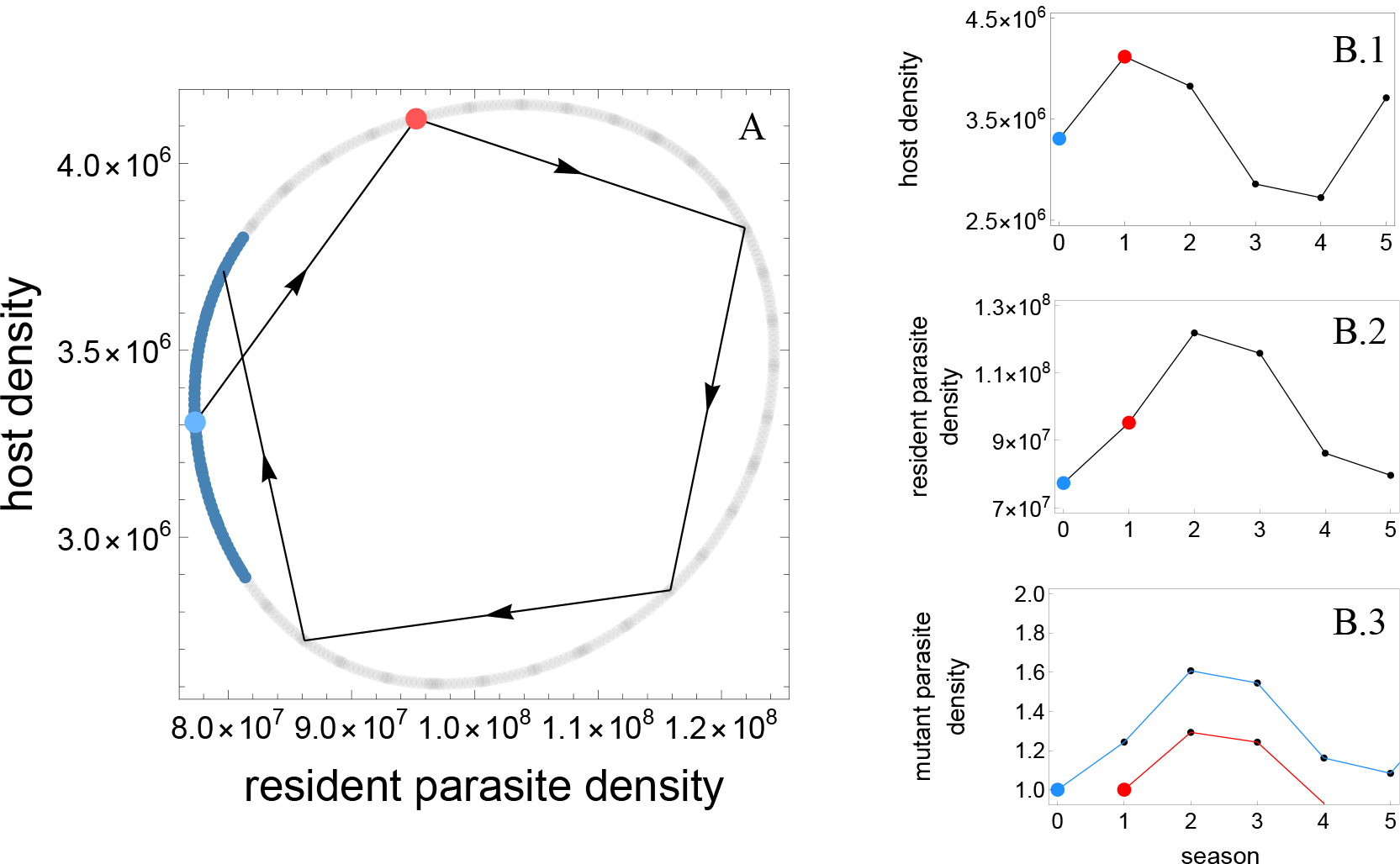
Mutant parasites with more adaptive virulence phenotypes often fail to invade when resident parasite and host dynamics are cycling. The phase plane (**A**) shows the discrete time limit cycle for host (*ŝ*) and resident parasite 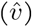 densities (*T* = 4; *t*_*l*_ = 1; *τ* = 2.8 in this example). The blue section denotes the phase of the parasite-host cycle when rare adaptive parasites can invade; the same mutant fails to invade when introduced at all other time points despite having the same selective advantage. The line (**A**) depicts the same iteration (6 seasons) of the quasi-periodic dynamics of this system as illustrated in **B.1** and **B.2**. An advantageous mutant fails to invade (red line, **B.3**) if introduced in seasons when host density will decrease (red point, **B.1**) and resident parasite density is moderate or high (red point, **B.2**). The same advantageous mutant can invade (blue line, **B.3**) and eventually replace the resident parasite if it is introduced when host density will increase (blue point, **B.1**) and resident parasite density is low (blue point, **B.2**). *T* = 4, *t*_*l*_ = 1, all other parameters found in Table 1.

## 4 Discussion

Host phenological patterns govern parasite densities directly through the timing and frequency of ecological interactions which can lead to an over-exploitation of hosts and subsequent parasite-host population cycles. Parasites can achieve sufficiently high densities in only some host phenological environments to destabilize the host-parasite dynamics that instigate quasi-periodic population cycles. The population cycles result from the classic consumer-resource ecological feedback where the parasite consumer over-exploits their host resource such that the host population cannot fully rebound in the following year. The resulting host population size is insufficient to support the now excessively-large parasite population which results in a dramatic decline in the number of parasites in the following year. The host population can then rebound due to the limited demographic impact of parasitism, thus allowing parasites to again over-exploit their hosts and restart the population cycle. Additionally, an evolutionary feedback can result from this consumer-resource ecological feedback. Parasite adaptation towards optimal trait values proceed more slowly when host-parasite dynamics are cycling. That is, many of the parasites with adaptive mutant phenotypes that arise in a cycling system will not increase in frequency and ultimately be lost from the population.

The observed parasite-host population cycles emerge from a delayed density dependent mechanism characteristic of consumer-resource feedbacks.^31^ In this system, the discrete host activity period introduces a delayed carryover effect in which the number of infected hosts in one season governs the host population size in the next. Although consumer-resource interactions can drive cycling in continuous time models, cycles are less likely to occur without an externally-imposed delay. The results of this study differ from those of prior studies describing consumer-resource feedbacks as this delayed density dependent mechanism causes population cycles only in phenological environments that support high parasite densities. Phenological patterns where hosts have shorter seasons and more synchronous emergence limit parasite deaths caused by environmental decay and infected-host deaths resulting in large parasite populations that can destabilize parasite-host dynamics and cause population cycles. By contrast, longer seasons and greater variation in emergence times among hosts support lower parasite densities which do not cause population cycles.

The stable parasite-host dynamics observed in some host phenological patterns challenges seminal theoretical studies demonstrating chaotic dynamics at all population growth rates of lethal parasites.^32^ Our results suggest that host phenology may stabilize host-parasite dynamics and explain why chaotic dynamics are not observed in all natural obligate-killer parasite systems. However, model parameters such as natural host mortality rate and parasite decay rate also modulate parasite population sizes and thus alter which phenological scenarios can lead to periodic population cycles (see Appendix C). Further, several factors that have been shown to impact the probability of dynamic population cycles not explored in this model could also modulate which phenological scenarios cycling could be expected.^33, 34^ For example, higher fecundity in infected hosts would likely stabilize the dynamics for a greater range of phenological patterns.

Population cycling resulting from a consumer-resource ecological feedback precipitates an eco-evolutionary feedback that affects the rate of adaptive evolution. Parasites with advantageous mutations always invade systems in which parasite densities are insufficient to destabilize parasite-host dynamics. That is, advantageous mutants displace residents both in systems where the parasite is not sufficiently adapted in a host phenological environment that could support high parasite densities and in systems where the host phenological pattern cannot support densities that cause population cycling even for the optimally-adapted parasite. By contrast, only a fraction of parasites with adaptive mutations introduced into cycling systems can invade, effectively reducing the rate of adaptive evolution. Parasites with adaptive mutations that do not invade fail to find a susceptible host when the resident parasite populations are large and the host population is small due to the numerical advantages of the resident. An assumption of the current model is that mutants are introduced at the beginning of random seasons, regardless of parasite population size. However, mutants are more likely to arise in large parasite populations in nature suggesting that the true impact of population cycling on the evolutionary rate is greater than estimated here.^35^

Our results suggest that variation in host phenology across geographic space could drive the observed differences in demographic dynamics. For example, parasite-host systems in more extreme latitudes and at higher altitudes are more likely to cycle than their conspecifics in less extreme environments.^36–38^ The activity periods in the more extreme environments tend to be shorter and hosts may emerge more synchronously,^39, 40^ in line with the predictions from the current model. These predictions could be tested empirically by studying the population dynamics of host-parasite systems in different geographic locations. Key parasite traits could also be measured to determine how parasite adaptation to different phenological patterns affected the differing demographic dynamics. Empirical data across locations could examine how phenology impacts species interactions and how that could cause differences in population densities, selection, and dynamical trajectories.

Several features of the current model can be altered to investigate more complex impacts of host phenology on parasite-host dynamics and eco-evolutionary feedbacks. For example, permitting host evolution in either parasite resistance or phenological patterns could drive additional eco-evolutionary feedbacks through changes in the strength of selection imposed on hosts by parasite infections. Future theoretical and empirical investigations into the impact of parasite-host cycles on the evolution of host resistance alleles, as seen in Gypsy moth populations,^41^ could determine if parasite-host co-evolution would stabilize population dynamics for a greater range of host phenological patterns. Similarly, the strength and possibly direction of selection pressures on hosts will fluctuate as the system cycles, potentially favoring alternative host phenological patterns that in turn select for parasite traits with lower impacts on host fitness. We will extend the current model to address these questions in future studies.

Relaxing some of the assumptions in this model is unlikely to qualitatively alter the major conclusions. For example, relaxing the monocylic parasite life cycle assumption will likely not change the result that cycles occur more readily in environments with short seasons and synchronous host emergence. Polycyclic parasites may even drive cycles for a larger range of phenological patterns as multiple infection cycles within a season can exacerbate decreases in host densities. In contrast, relaxing the obligate-killer assumption will likely decrease the range of phenological patterns that can drive cycles by decreasing the impact on host fitness. Although the model as presented applies to only a narrow range of parasites in nature, many more parasite-host systems conform to models that include these extensions such as soil-borne plant pathogens, demicyclic rusts, post-harvest diseases, and many diseases systems infecting univoltine insects.^42–45^

Environmental conditions such as phenology impact the frequency of inter-species interactions and thus the ecological importance of the interaction on population demography. Here we show that short host seasons and synchronous host emergence can allow parasites to reach densities that destabilize population dynamics and cause demographic cycling. The rate of adaptive parasite evolution in a cycling population is substantially slower than in an equilibrial population as beneficial mutations are likely to go extinct when host population sizes are small or parasite population sizes are large. These results demonstrate that externally-imposed environmental conditions such as host phenology can be important determinants of cycling. Further, it is important to consider ecological dynamics when predicting evolution by natural selection.

## Appendix A

In Appendix A we find analytical solutions for equations 1a-c from the main text to study parasite fitness given different host phenological patterns.

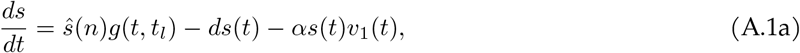

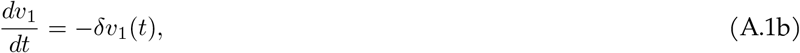

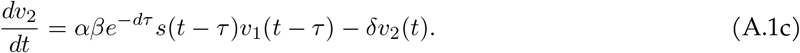

with initial conditions: 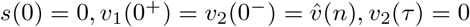.

(A.1a-c) is solved analytically by describing host emergence using a uniform distribution

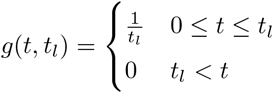

To solve the dynamics during the host’s activity period, we first find the analytical solution for *v*_1_(*t*):

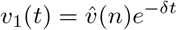

We then use *v*_1_(*t*) to find the time-dependent solutions for both *s*(*t*) and *v*_2_(*t*):

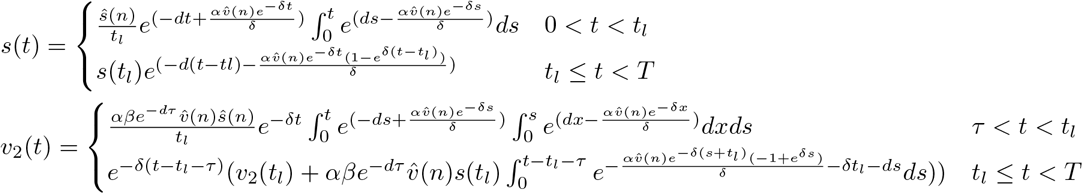

where *s*(*t*_*l*_) and *v*_2_(*t*_*l*_) are the densities of *s* and *v*_2_ when the emergence period of *s* ends.

Within-season dynamics are coupled to recurrence equations that describe host and parasite between-season dynamics. The total population of new parasites at the end of the season, *v*_2_(*T*), gives rise to next season’s starting parasite population, *i*.*e*.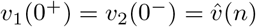. A parasite introduced into a naive host population persists or goes extinct depending on the length of the host emergence period and season length. The stability of the parasite-free equilibrium is determined by the production of *v*_2_ resulting from infection of *s* given by

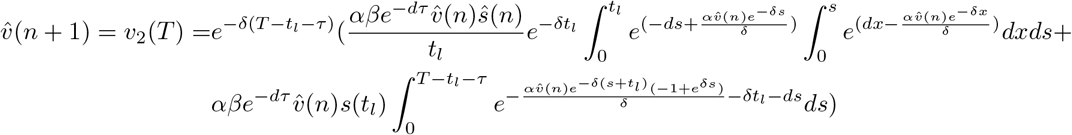

The parasite-free equilibrium is unstable and a single parasite introduced into the system at the beginning of the season will persist if the density of *v*_2_ produced by time *T* is greater than or equal to 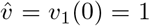 (*i*.*e. v*_2_(*T*) *≥* 1, modulus is greater than unity). This expression is a measure of a parasite’s fitness when rare given different host phenological patterns.

The total population of uninfected hosts at the end of the season, *s*(*T*), reproduce and give rise to next season’s host cohort, given by the map

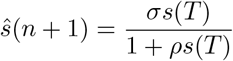

where *σ* is host fecundity and *ρ* is the density dependent parameter.

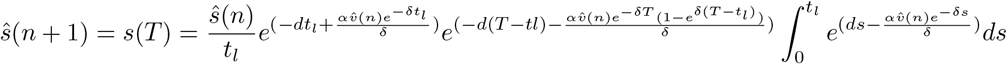

We can find the values of 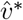 and *ŝ∗* numerically that satisfy

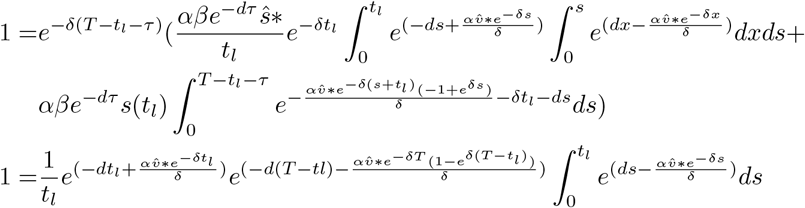

If we define 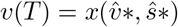 and 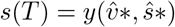, we can write the Jacobian for the between-season stability analysis as

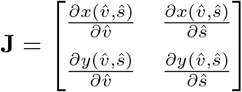

We cannot solve this model further analytically as the partial derivatives w.r.t. 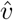 are transcendental. We are however able to find the eigenvalues of this Jacobian numerically. The leading eigenvalue is complex conjugate. The real part of the leading eigenvalue is less than one when the system dynamics are stable and greater than 1 when the system dynamics are cycling. This suggests that a Neimark-Sacker bifurcation drives the system to cycle.^46^

## Appendix B

In Appendix B we find analytical solutions for equations 2a-e from the main text to study the evolution of parasite virulence given different host phenological patterns. Note that we primarily used numerical simulations in the main text to determine the outcome of parasite evolution as this analytical solution only holds when host and resident parasite populations are at a stable equilibrium when the mutant is introduced.

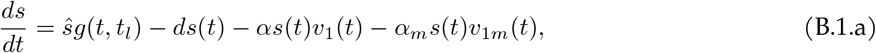

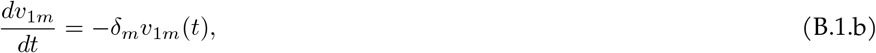

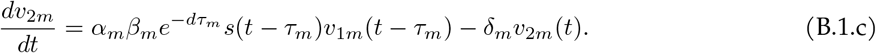

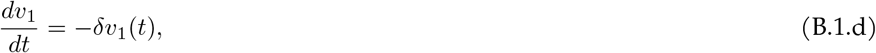

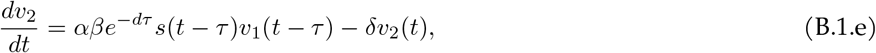

with initial conditions: *s*(0) = 0, *v*_1*m*_(0^+^) = *v*_2*m*_(0^*-*^), *v*_2*m*_(*τ*) = 0, *v*_1_(0^+^) = *v*_2_(0^*-*^), *v*_2_(*τ*) = 0. *m* subscripts refer to the invading mutant parasite and its corresponding traits.

(B.1.a-c) has the following time-dependent solution:

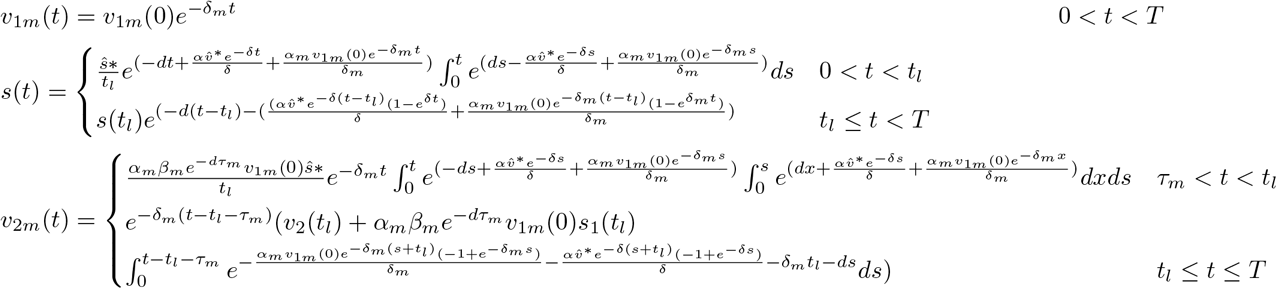

The invasion fitness of a rare mutant parasite is given by the density of *v*_2*m*_ produced by the end of the season. The mutant parasite invades in a given host phenological scenario if the density of *v*_2*m*_ produced by time *T* is greater than or equal to the initial *v*_1*m*_(0) = 1 introduced at the start of the season (*v*_2*m*_(*T*) *≥* 1).

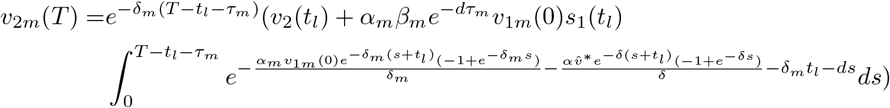

We can use *v*_2*m*_(*T*) to find optimal virulence for a given host phenological scenario by finding the trait value that maximizes *v*_2*m*_(*T*). That is, the virulence trait, *τ* ^*∗*^, that satisfies

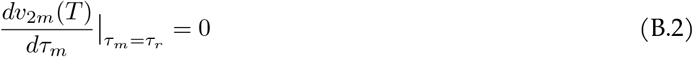

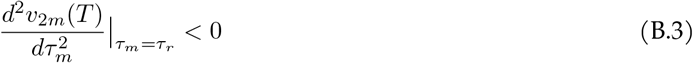

We primarily used numerical simulation to determine parasite evolutionary endpoints as the analytical solution does not reliably predict the invasion fitness of rare mutant parasites invading populations with cycling dynamics.

